# Developmental regulation of Drosophila dosage compensation in a 3D genome context

**DOI:** 10.1101/2025.02.28.640876

**Authors:** Lubna Younas, Mujahid Ali, Xinpei Zhang, Catherine Regnard, Qi Zhou

## Abstract

Chromosomal dosage compensation (DC) restores the expression balance of X chromosome between sexes, and is realised in Drosophila by the coordination of enriched X-linked PionX, HAS sequence elements and 1.688^3F^ satellites, resulting in doubling the transcription of X-linked genes in males. We hypothesize that DC must be finely tuned during development and ask what are the underlying molecular mechanisms in this work. To inspect this, we collect new and published histone modification and 3D genome data of sexed samples from embryos to adults of Drosophila. We find that at least 30% of the Drosophila genes are bound by the DC complex or characteristic H4K16ac modifications specifically in only one or more tissues, and they show a higher H3K27me3 binding level than the rest DC genes constitutively compensated throughout development. The chrX harbors developmentally increasing long-range contacts (LRCs) relative to autosomes, and specifically in males. Integrative analyses with DC genes and regulatory sequences show that HAS and housekeeping genes preferentially located at the chromatin domain boundaries form stable LRCs. While PionXs and 1.688^3F^ only form LRCs with the developmentally regulated DC genes in embryos and testes. Our results indicate that the Drosophila male chrX is characterised by two types of stable and developmentally regulated LRCs coordinating the dosage compensation to the different functional background of genes.

## Introduction

Sex chromosomes often have evolved dramatically different genomic and epigenomic compositions: the Y chromosome has usually lost most functional genes and become heterochromatic due to lack of homologous recombination, while the male X chromosome (chrX) stays largely euchromatic (Bachtrog, 2013). This renders an imbalanced gene dosage between the X and autosomes and sometimes selects for evolution of diverse mechanisms of chromosome wide dosage compensation (DC)(Ohno, 1966; Charlesworth, 1978; Gu and Walters, 2017). In genetic model organisms, DC is a fundamental process that has been demonstrated to be critical for the viability and normal development(Disteche, 2012; Copur *et al*., 2018). It equalizes the expression level between sexes, through either upregulating the male X (e.g., in Drosophila(Conrad and Akhtar, 2012; Lucchesi and Kuroda, 2015) and green anole lizard(Marin *et al*., 2017)), or downregulating the female chrX(e.g., in eutherian mammals(Heard and Disteche, 2006)) in transcription or translation (e.g., recently reported in chicken and platypus(Lister *et al*., 2024)). The initiation, targeting and spreading processes of DC have been the forefront research paradigms of revealing mechanisms of transcriptional regulation that involve dynamic coordination between the local *cis*- and *trans*-regulatory elements, in the context of global modulation of 3D chromatin architecture of an entire chromosome, with *D. melanogaster* having one of the most extensively studied DC mechanisms.

In Drosophila, the current model proposes that the male-specific lethal (MSL) complex recognizes the X-linked high-affinity binding sites (HAS(Straub *et al*., 2008), or chromatin entry sites, CESsHA (Alekseyenko *et al*., 2008)) mediated by the *roX* RNAs(Meller and Rattner, 2002). The MSL complex then deposits the active histone post-translational modification mark (HPTM) H4K16ac (histone H4 acetylation at lysine 16(Gelbart *et al*., 2009)) onto or nearby the transcribed X-linked genes enriched for another active HPTM H3K36me3(Larschan *et al*., 2007). This produces an overall hyperactive chromatin configuration and doubled transcription output of the male chrX. Many details of how the MSL complex specifically targets the chrX remain to be understood, but it probably first binds to a subset of HASs, termed pioneering sites on the X (PionX) during its *de novo* assembly. The recognition process requires synergistic action of the maternally deposited protein CLAMP(Rieder, Jordan and Larschan, 2019; Jordan and Larschan, 2021), and is promoted by small interfering RNAs encoded by a satellite repeat cluster 1.688^3F^ located at the 3F cytogenetic region of chrX(Menon *et al*., 2014). All of the three critical sequence elements, HASs, PionXs and 1.688^3F^ are enriched on the Drosophila chrX relative to autosomes, but to a different degree, and HASs and PionXs are enriched for different variants of 21bp GA-rich motifs called MRE (MSL recognition elements(Alekseyenko *et al*., 2008)).

Similar to mammals and nematodes (reviewed in (Jordan, Rieder and Larschan, 2019)), in Drosophila the subsequent spreading of MSL complex is facilitated by or has consequential impacts on the distinctive chromatin architecture of the chrX. The male chrX is more accessible, and exhibits more mid-/long-range contacts compared to autosomes(Pal *et al*., 2019), and the MSL complex could have taken advantage of the spatial proximity between HASs and PionXs to effectively spread to nearby genes (Ramírez *et al*., 2015; Schauer *et al*., 2017). Besides these abundant mechanistic studies, variations of DC and its targeted genes during the developmental process and in different cellular contexts remain largely unknown, as most previous studies in Drosophila were confined to cell lines or embryos.

There must be strong natural selection against the DC instinctively doubling the transcription level of every X-linked gene in each type of male cell, given the tremendous diversities of dosage sensitivity (thus selective pressures for evolving DC) and tissue-specific functions between individual genes, and the heterogeneous local chromatin environment of chrX. Indeed, only a subset of X-linked genes are subjected to non-canonical incomplete DC before the blastoderm stage and in male germ cells(Lott *et al*., 2011; Li *et al*., 2021; Witt *et al*., 2021), where the Drosophila MSL-mediated DC is absent. In humans, over 15% of X-linked genes escape DC or X-inactivation(Carrel and Willard, 2005), and 5.8% of the genes escape X-inactivation in one of the studied tissues(Tukiainen *et al*., 2017). Transcriptome evidence for tissue-dependent DC has also been reported in mice(Berletch *et al*., 2015), *D. pseudoobscura* (Nozawa *et al*., 2014), with the precise regulatory mechanisms of escaping the DC unclear(Picard *et al*., 2019). However, transcriptome differences between sexes, as a common practice of defining presence/absence of DC on individual genes, is not necessarily an accurate reflection of DC, because it is impacted by the opposite forces between DC and sexual selection(Mullon *et al*., 2015; Huylmans, Macon and Vicoso, 2017).

To overcome this confounding factor, and more importantly uncover the patterns and mechanisms of developmental regulation of DC in *D. melanogaster*, here we collect chromatin immunoprecipitation sequencing (ChIP-seq) data targeting the characteristic DC protein (MSL2) or HPTM (H4K16ac) spanning major developmental stages and tissue types, in order to unambiguously define the DC status of each X-linked gene in each sample by the significant MSL2/H4K16ac occupancy, rather the transcription level. With other active or repressive HPTM ChIP-seq (e.g., the polycomb and heterochromatin marks H3K27me3 and H3K9me2/3) and newly collected high throughput chromatin conformation capture (Hi-C) data in the corresponding stage/tissue of Drosophila (**Figure 1**), we seek to answer: 1) How variable is the MSL-mediated DC across development and what are the epigenetic signatures that distinguish the genes that are unanimously dosage compensated (constitutive dosage compensated genes, CDC genes), those that are compensated in some but not all tissues (facultative DC genes, FDC genes) and those that are compensated in only one tissue or stage (specific DC, SDC genes)? 2) How variable is the 3D chromatin architecture of the chrX, under the influence of developmentally regulated DC, across tissues and stages? And how are different categories of DC genes synergistically regulated by such chromatin contacts, in addition to the local chromatin configuration found in 1)? 3) What are the different roles of three known X-linked sequence elements PionX, HAS and 1.688^3F^ in regulating different categories of DC genes? Our results provide novel insights into how DC is dynamically regulated in the different contexts of genes, chromatin and cell types, and have broad implications on other DC systems.

**Figure 1.**
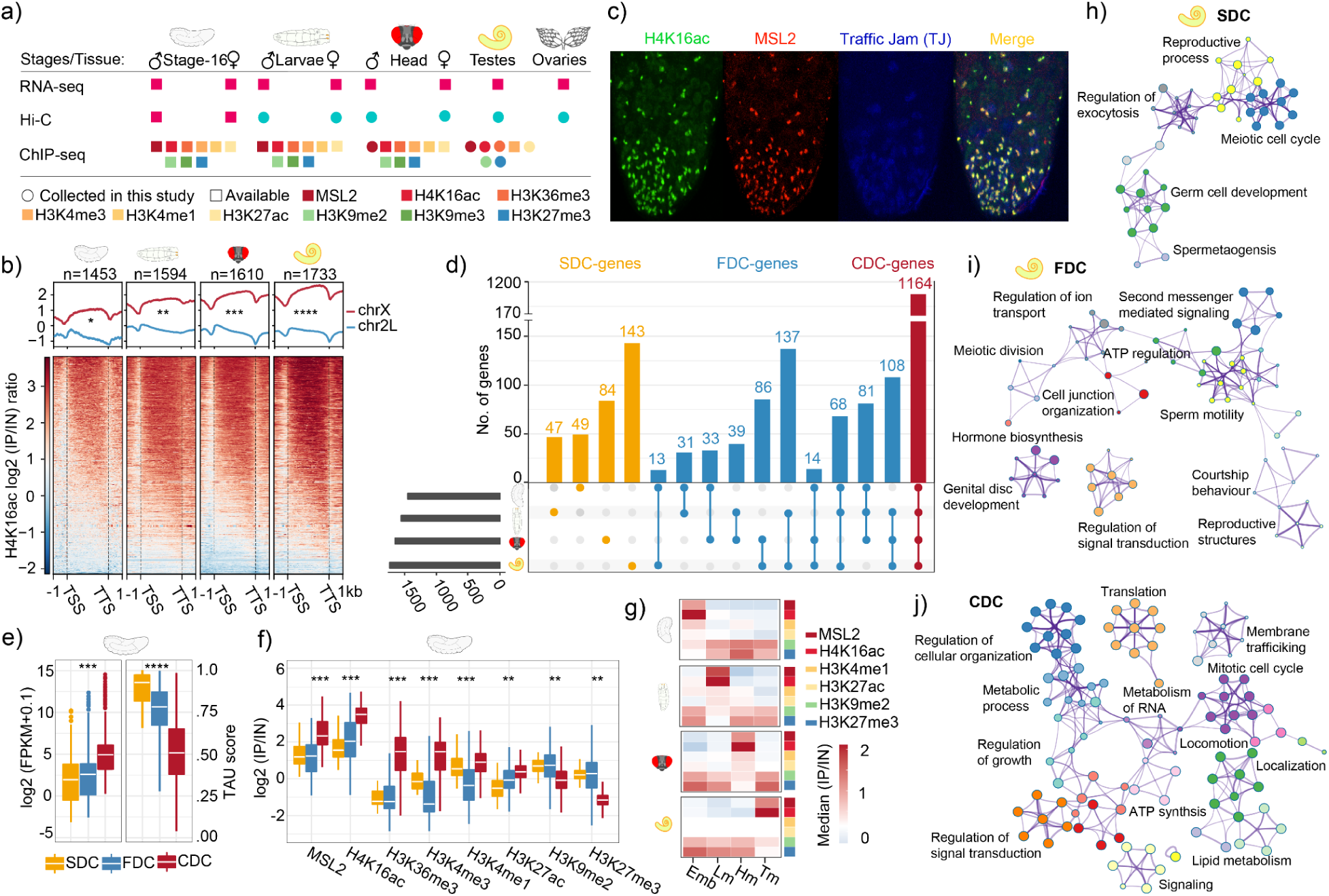
Developmental variations of dosage compensated genes and their epigenetic states. **(a)** Schematic representation of the datasets and developmental stages used in the study. Sex-specific Hi-C, ChIP-seq, and RNA-seq data were collected from stage-16 embryos, 3rd instar larvae, adult heads, and testes. Colored circles indicate the type of chromatin mark analyzed with ChIP-seq, with new data collected in this study denoted by outlined circles. **(b)** Differential enrichment of H4K16ac and MSL2 on chromosome X (chrX) compared to autosomes (chr2L) across development. Top panel: Box plots showing normalized binding strengths on chrX versus autosomes, with statistically significant differences marked (**P < 0.01, ***P < 0.001, ****P < 0.0001, Wilcoxon test). Bottom panel: Heatmaps depict the log2 ratio of IP (immunoprecipitation) over IN (input) signals, revealing the distribution of H4K16ac and MSL2 across chrX and chr2L at different developmental stages. **(c)** Immunofluorescence staining of D. *melanogaster* testes for H4K16ac (green), MSL2 (red), and Traffic Jam (TJ, blue). Colocalization in the merged image indicates the presence of canonical dosage compensation within the somatic cells of the testes. **(d)** Quantification of specific DC (SDC), facultative DC (FDC), and constitutive DC (CDC) genes across developmental stages and tissues, represented as the number of genes in each category. **(e)** Box plots illustrating the variation in gene expression levels (FPKM) and TAU scores among SDC, FDC, and CDC genes across developmental stages and tissues (***P < 0.001, ****P < 0.0001, Wilcoxon test). **(f)** IP/IN ratios of histone post-translational modifications (HPTMs) at SDC, FDC, and CDC genes. Significant differences marked (**P < 0.01, ***P < 0.001, Wilcoxon test). **(g)** Median IP/IN ratios of HPTMs at SDC genes across different stages and tissues, displayed as heatmaps. The color gradient represents the median log2 ratio of IP over IN, with red indicating higher enrichment. **(h), (i), (j)** Network plots of Gene Ontology (GO) terms associated with SDC, FDC, and CDC genes, respectively. Nodes represent enriched biological processes, and edges denote shared genes between GO terms. The nodes’ size correlates with the enrichment significance (adjusted P < 0.05, Fisher’s Exact test).

## Results

### Developmental variations of dosage compensated genes and their epigenetic states

We use the published sexed transcriptome and ChIP-seq data of stage 16 embryo (Renschler *et al*., 2019), third instar larvae (Valsecchi *et al*., 2018), virgin adult heads(modENCODE Consortium *et al*., 2010) targeting the DC protein or characteristic HPTM (MSL2, H4K16ac), and other active (H3K36me3, H3K4me1/3, H3K27ac) and repressive (H3K9me2/3, H3K27me3) HPTMs. In this work we further collect 6 matched ChIP-seq data from testis, and sexed Hi-C data for larvae, heads and gonads (**Figure 1a**). As expected, the normalized binding strengths (ChIP vs. input, IP/IN ratio) of H4K16ac (**Figure 1b**) and MSL2 (**Supplementary Fig. 1a-b**) are significantly (*P* < 0.05, Wilcoxon test) higher on the X-linked vs. autosomal genes. And H4K16ac exhibits a reported bias toward 3’ gene body on X-linked genes, while autosomal genes have an opposite bias pattern(Gelbart *et al*., 2009) (**Figure 1b**). The X vs. autosome differences of H4K16ac are not significant until after the zygotic genome activation (embryonic stage 7(Samata *et al*., 2020)) and become gradually more pronounced during development into the adult, with testis showing the highest binding strength of H4K16ac on X-linked genes among the studied samples (**Supplementary Fig. 1c**). By the distinctive chromosomal distributions of H4K16ac/MSL2 binding strength, we are able to define a respective cutoff in each tissue/stage that separates the X and autosomes for defining the DC gene as bound by either H4K16ac or MSL2 (**Supplementary Fig. 1d-e**). There is indeed gradual establishment of DC, particularly at the embryonic stages (**Supplementary Fig.1f-g**), indicated by the increasing number of DC genes from 668 to 1733 by development (**Supplementary Fig. 1c**). Previous studies suggested MSL-mediated DC is absent in male germ cells(Rastelli and Kuroda, 1998; Witt *et al*., 2021), and we confirm this here by immunofluorescence (IF) staining: antibodies against H4K16ac and MSL2 only stain the cyst cells but not germ cells (**Figure 1c**). This indicates that our H4K16ac/MSL2 ChIP-seq data from testis only reflect patterns from cyst cells.

To inspect the developmental variation of DC, we examine the overlap of genes and identify 1164 (62% of total X-linked genes) CDC genes, 426 (23%) FDC genes, and 143 SDC (8%) genes, with testis consistently showing the highest number of FDC or SDC genes (**Figure 1d**). These three types of DC genes show distinctive expression and epigenetic features. CDC genes have significantly (*P* < 0.05, Wilcoxon test) higher transcription levels in all examined tissues/stages than the other two sets of DC genes, but a significantly (*P* < 0.05, Wilcoxon test) lower level of tissue specificity, indicating that CDC gene are enriched for housekeeping genes (**Figure 1e, Supplementary Fig. 1h-i**). The transcription pattern is corroborated by the ChIP-seq data that CDC genes have significantly (*P* < 0.05, Wilcoxon test) higher normalized binding strengths of active HPTMs including H4K16ac and that of MSL2, but lower (*P* < 0.05, Wilcoxon test) binding strengths of repressive HPTMs than FDC and SDC genes (**Figure 1f, Supplementary Fig. 2a**). In order to understand the epigenetic mechanisms regulating the SDC genes, we compare the normalized HPTM binding strengths of these genes across the studied samples. As expected, SDC genes show specific bindings of MSL2/H4K16ac in the tissue/stage where they are compensated, and also significantly (*P* < 0.05, t-test) higher binding strengths of other active HPTMs (e.g., H3K4me1/3, H3K27ac) than other tissues where they are not compensated. But at the same time in all of the other tissues/stages they are also enriched for repressive HPTMs H3K9me2 and H3K27me3, i.e., silenced for transcription (**Figure 1g, Supplementary Fig. 2b**). It is important to note that the genes nearby SDC genes do not show a similar epigenetic feature across tissues/stages and are mainly CDC genes, suggesting a rather restricted chromatin domain regulating these SDC genes (**Supplementary Fig. 2c**).

These specific genes show highly relevant functional enrichment to the respective tissue or stage by our gene ontology (GO) analyses (**Supplementary Fig. 2d-i**). For example, the SDC and FDC genes in testis are enriched (*P* < 0.05, Fisher’s exact test) for GOs of ‘sperm motility’, ‘germ cell development’, ‘spermatogenesis’ (**Figure 1h, i**), and enriched (*P* < 0.05, ** test) for mutant phenotypes involving cyst cell structures of spectrosome and fusome, or those showing defects in meiosis and fertility (**Supplementary Fig. 3a, h**). These testis SDC/FDC genes are ‘overcompensated’ and enriched (*P* < 0.05, Fisher’s exact test) for male-biased genes relative to other X-linked genes (**Supplementary Fig. 4a-c**). Those in male heads are enriched for GOs of ‘courtship behavior’ (e.g., the FDC gene *yellow(Massey et al., 2019)*), ‘neural projections’ etc. (**Supplementary Fig. 2f, i**), and enriched for mutant phenotypes of defects in circadian rhythms (e.g., the SDC gene *period(Lee, Bae and Edery, 1999)*) or mushroom body abnormalities (**Supplementary Fig. 3c, f**). By contrast CDC genes, mostly housekeeping genes, are enriched (*P* < 0.05, Fisher’s exact test) for female-biased genes, and GOs of ‘metabolism of RNA’, ‘translation’, ‘regulation of growth’ etc (**Figure 1j, Supplementary Fig. 4d-f**). In summary, we estimate that over 30% of the X-linked genes are not dosage compensated in all the stages or tissues, probably because their specific functions in one tissue or stage select for absence of DC and epigenetic silencing in all the other stages and tissues.

### Male chrX harbors developmentally increased long-range contacts

It was previously reported that there are long-range contacts (LRC) specific to the male chrX by studying the Drosophila embryos and cell lines(Pal *et al*., 2019). We ask here whether there is a developmental increase of LRCs, and whether it associates with that of the DC genes found above (**Figure 1b, Supplementary Fig. 1c**). During development, we find the chromatin interaction strengths between the homotypic active (A-A) or inactive (B-B) compartments in males but not in females become significantly (*P* < 0.05, Wilcoxon test) stronger between consecutive stages, and the (**Supplementary Fig. 5a-c**) male chrX always exhibits stronger (*P* < 0.05, Wilcoxon test) contacts than autosomes (**Figure 2a, Supplementary Fig. 5b**). Consistently, there is a significant (*P* < 0.05, Wilcoxon test) increase of male vs. female ratio of Hi-C contact signals on the chrX but not on autosomes during development (**Figure 2b, Supplementary Fig. 5d**). As the chromatin compartment length is estimated to be on average 390kb long by Hi-C data of this work. We test the hypothesis that these male-specific chrX contacts spanning different compartments are mainly attributed to the LRC rather than the short-range contacts (SRC) (**Supplementary Fig. 5e**). Indeed, analyses of distance-dependent contact frequency decay on each chromosome show that when the distance between any of the two contacting genomic windows is higher than 1Mb, the contact frequency of the male but not the female chrX has a clear shift away from those of autosomes toward a higher level (**Figure 2c**). In particular, the LRC difference between the male chrX vs. autosomes also becomes more significant (*P* < 0.05, Wilcoxon test) by development. We examine the LRC regions that only appear in the later stage than the previous stage and find that SDC or FDC genes predominantly (between 88% to 96% across samples of the LRC regions (**Supplementary Fig. 5f**) overlapped with these regions. Similarly, the SDC/FDC genes bound by H3K27me3 form specific or facultative LRCs (**Figure 2d**) distinctive from the stable LRCs. These results together suggest that the SDC or FDC genes form specific or facultative LRCs distinctive from the stable LRCs shared across stage/tissue on the male chrX, that are by contrast enriched between CDC genes (**Supplementary Fig. 5g**).

**Figure 2.**
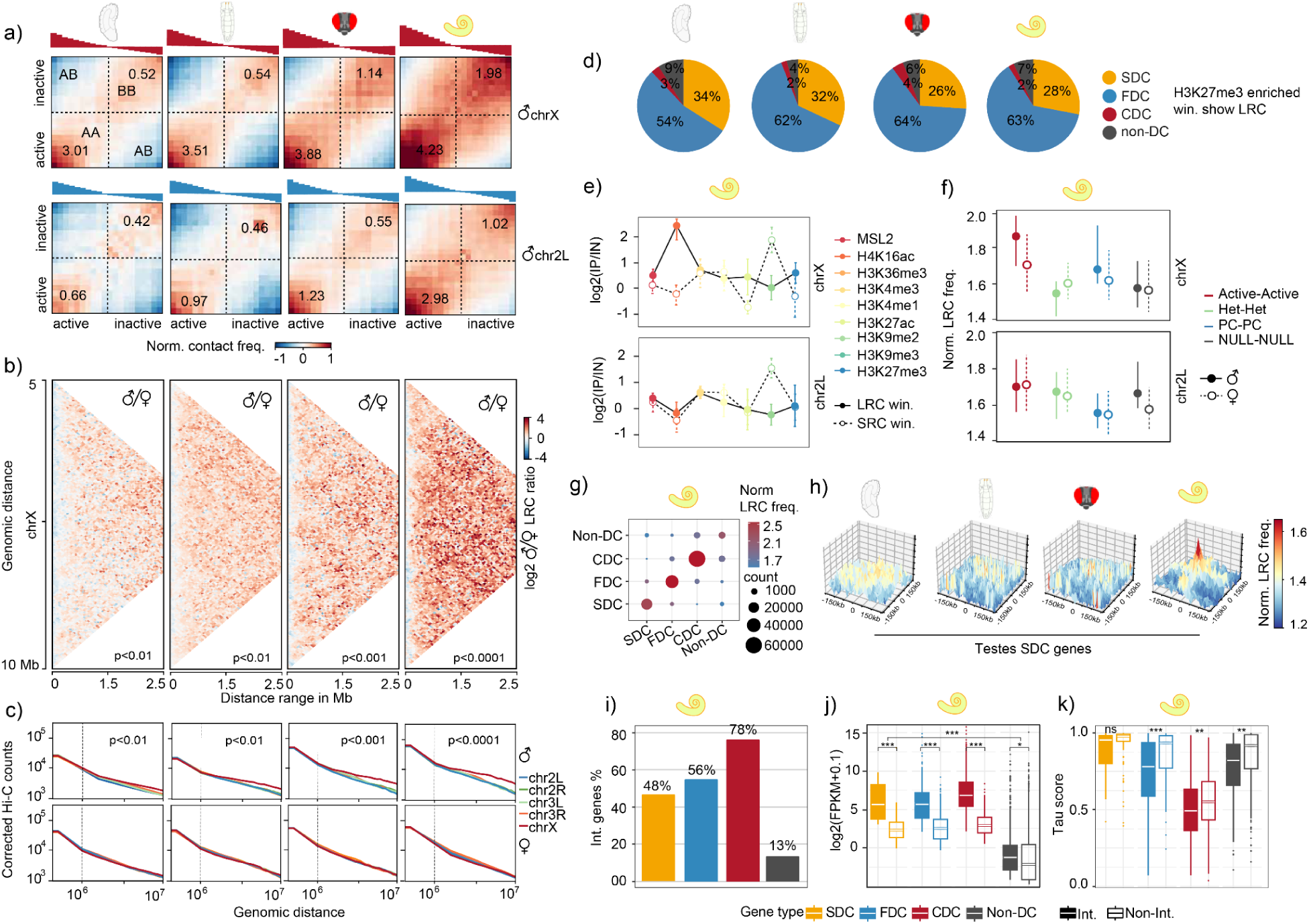
Male chrX harbors developmentally increased long-range contacts. **(a)** Chromosome interaction matrix comparing active (A) and inactive (B) compartment contacts across chrX and chr2L. Numbers indicate log2 observed/expected (O/E) interaction frequencies, with higher values in active compartments on dosage compensated male chrX than autosomal chr2L. **(b)** Sex-specific Hi-C contact matrices for chrX within a 0 to 3Mb distance range, highlighting the increased long-range contacts (500 kb - 3Mb) in males, represented by the red to blue gradient denoting higher to lower male/female Hi-C signal ratio, respectively. **(c)** Log-log plot of distance-dependent interaction frequency decay for chromosomal arms 2L, 3R, 3L, and X, showing chrX deviating from autosomal arms in males with higher frequency LRCs. The dotted line represents the cutoff between SRCs (<1Mb distance range) and LRCs (>1Mb distance range). **(d)** H3K27me3 enriched bins that also participated in the LRCs. **(e)** Enrichment of histone post-translational modifications (HPTMs) across short- and long-range interacting bins on chrX and chr2L. Notably, H4K16ac displays significantly higher enrichment in long-range contacts on chrX (P < 0.001, Wilcoxon test). **(f)** A comparison of normalized LRC frequencies between pairs of long-range interacting bins categorized by their chromatin states demonstrates that active chromatin domains have significantly higher contacts on male chrX. **(g)** Analysis of interaction patterns among dosage compensation gene categories (CDC, FDC, Non-DC) on chrX, showing a higher frequency of contacts among CDC genes compared to FDC and SDC genes. **(h)** Three-dimensional interaction landscapes display the normalized LRC frequencies between testes-specific DC (SDC) genes across development. **(i)** Percentage of genes within each DC category (CDC, FDC, Non-DC) that participate in long-range contacts, with CDC genes more frequently engaged. **(j)** Box plot comparing expression levels (FPKM) of genes involved in long-range contacts to those not involved, indicating higher expression for interacting genes. **(k)** TAU score analysis of interacting versus non-interacting genes, revealing that genes with long-range contacts have lower tissue specificity, reflected by lower TAU scores. This figure provides a multi-dimensional view of how dosage compensation in D. melanogaster influences the 3D chromatin architecture, particularly enhancing long-range contacts on the male chrX. The results underscore the unique role of H4K16ac in facilitating these long-range contacts, potentially shaping gene expression during development.

The epigenomic features of these male X-linked LRC regions are consistent with their overlaps with the SDC/FDC genes (**Figure 1f**). We find that the X-linked LRC regions are characterized by a significantly (*P* < 0.05, Wilcoxon test) stronger bindings of H4K16ac and polycomb mark H3K27me3, but not MSL2, than the SRC regions in males but not in females. And the two HPTMs’ normalised binding levels also significantly increase (*P* < 0.05, Wilcoxon test) by the development course (**Figure 2e, Supplementary Fig. 6a-b**). *Vice versa*, the X-linked regions decorated by active HPTMs or H3K27me3 show a significantly (*P* < 0.05, Wilcoxon test) stronger LRC strength in male than in female and by development (*P* < 0.05, Wilcoxon test) (**Figure 2f, Supplementary Fig. 6c-d**). This suggests that the X-linked LRC regions in males are likely associated with the subsequent spreading rather than the original establishment of DC, as MSL2 marks the primary binding sites of the DC complex. When further examining the LRCs in a pairwise manner, we find that most of them are between DC genes of the same type, e.g., between CDC genes rather than between CDC and SDC or FDC genes (**Figure 2g, Supplementary Fig. 6e**). The epigenomic features of SDC genes align with their overlap with specific or facultative LRCs (**Figure 2d**). Genes compensated in one stage or tissue exhibit significantly higher LRC frequencies in that tissue. While, these interactions are weaker or absent in other stages or tissues where the gene is not compensated (**Figure 2h, Supplementary Fig. 6f**). A much higher percentage of CDC genes (62%) is found to show pairwise LRCs than that of FDC or SDC genes (56% or 48%) (**Figure 2i, Supplementary Fig. 7a-b**). All X-linked genes with pairwise LRCs, including all types of dosage compensated genes and a few genes that are not compensated, show a significantly higher transcription level, and lower tissue expression specificity (hence broader tissue expression pattern), than other DC genes without pairwise LRCs (**Figure 2j-k, Supplementary Fig. 7c**). In addition, if we study SDC/FDC genes without LRCs, we do not find enrichment of GO terms as relevant as those with LRCs with the specific tissues (**Supplementary Fig. 7d-g**). This result suggests that LRCs are associated with regulation of genes with tissue specific functions.

### Different genes and regulatory sequences of dosage compensation have respective long range contacts

Three classes of regulatory sequence elements enriched on the chrX relative to autosomes, PionX(Villa *et al*., 2016), HAS (Alekseyenko *et al*., 2008) and 1.688^3F^ (Menon *et al*., 2014) are known to initiate, attract or promote the recruitment of MSL complex along the chrX. We next ask whether and how they are engaged in the LRCs that were shown above to be a characteristic and developmental feature of chrX. We find that all three classes of elements, with some developmental variations, exhibit significantly (Wilcoxon test, *P* < 0.05) stronger pairwise LRC frequencies than SRCs in males (**Supplementary Fig. 8a**), and LRCs between these elements are also (Wilcoxon test, *P* < 0.05) stronger than those in females (**Supplementary Fig. 8b**). There are also cross interactions of LRCs between PionX and HAS, but not between 1.688^3F^ satellites and PionX/HAS elements (**Supplementary Fig. 8c**). Particularly, the LRC frequency between HASs, but not PinoXs and 1.688^3F^ satellites, show a similar increase (Wilcoxon test, *P* < 0.05 between consecutive stages) during development with that of the entire chrX (**Figure 2**). This leads to the intriguing hypothesis to be tested in future that HASs might mediate the LRCs during development. While PionX elements and 1.688^3F^ repeats show much weaker, or no LRCs in the larvae and head tissues (**Figure 3a**).

**Figure. 3:**
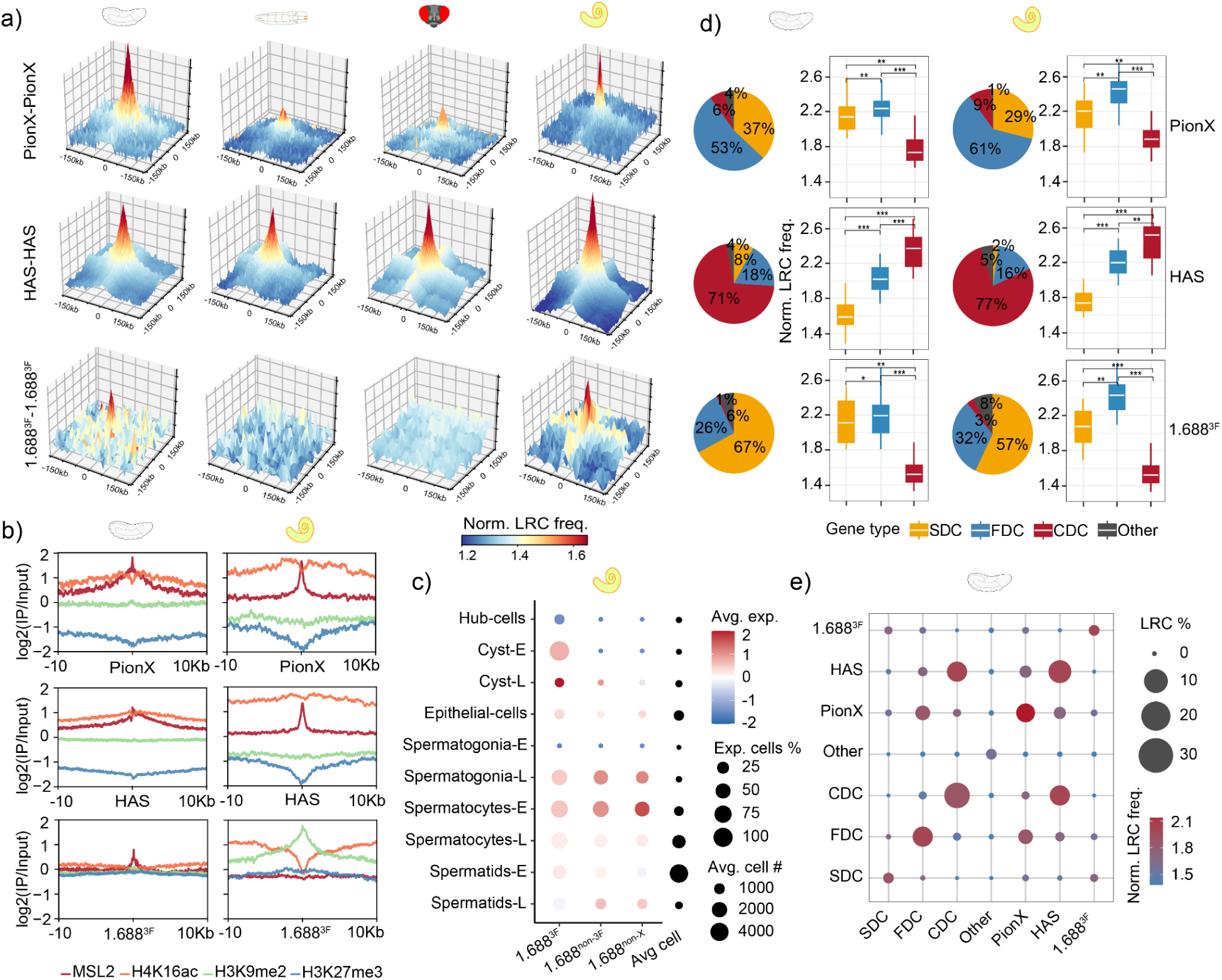
Different genes and regulatory sequences of dosage compensation have respective long range contacts. **(a)** Three-dimensional interaction landscapes display the normalized LRC frequency between pairs of regulatory elements (PionX-PionX, HAS-HAS and 1.688^3F^-1.688^3F^) over 1Mb distance range. These patterns highlight the distinct interaction behavior of these elements in embryos and testes compared to other developmental stages. The color bar represents the LRC frequencies lower (blue) to higher (red). **(b)** Enrichment profiles for MSL2, H4K16ac, H3K9me2, and H3K27me3 around pioneering sites on X (PionX), high-affinity sites (HAS), and the 1.688^3F^ satellite repeat along the dosage compensated chrX. These profiles illustrate the dynamic binding patterns of these elements in the embryo (first column) and testes (second column). **(c)** Transcriptional activity of 1.688 satellites with the published single-cell RNA-seq data of D. *melanogaster* adult testis. Circle size represents the percentage of expressed cells within each cell type, whereas blue to red color represents the average expression lower to higher respectively. **(d)** Pie charts on the right, represent the percentage distribution of different gene classes (including the flanking 5kb regions) around PionX, HAS and 1.688^3F^ satellites. The boxplot on left represents the normalised LRC frequencies between the PionXs, HAS or the 1.688^3F^ satellites within the SDC, FDC and CDC genes. Statistical significance was calculated using (Wilcoxon test, *P < 0.05, **P < 0.005, ***P < 0.0005). **(e)** The dot plot represents the fraction of chrX pair-wise LRCs involving different DC gene categories (SDC, FDC, CDC and non-DC) and *cis*-elements (PionX, HAS, and 1.688^3F^). The circle size represents the percentage distribution of total LRCs within and between different elements in the embryo. The color scale represents the normalized LRC frequency lower to higher as blue to red respectively.

Nevertheless, the LRCs between the 1.688^3F^ satellites in embryos and testes are of particular interest, because a recent genetic study suggested the satellites contribute to establishment of DC in embryos in parallel to HASs/PionXs (Makki and Meller, 2024). Moreover, 1.688 satellites elsewhere on the chrX (1.688^non-3F^), including those reported in the cytogenetic regions 1A and 3C of chrX (Menon *et al*., 2014) and autosomes (1.688^non-X^), do not show such pronounced LRCs (**Supplementary Fig. 9a-b**), with our stringent bioinformatic procedure to avoid cross-mapping between different clusters of 1.688 satellite repeats (**Materials and Methods**). It was shown in larvae that the 1.688^3F^ satellites do not directly bind to the MSL complex (Menon *et al*., 2014). However, in this work we find specifically at the embryonic stage an enrichment of MSL2, but not H4K16ac at the 1.688^3F^ satellites (**Figure 3b**). Other 1.688 satellites show a similar, but significantly (Wilcoxon test, *P* < 0.05) weaker levels of enrichment pattern (**Supplementary Fig. 9c**). This provides new evidence that 1.688^3F^ satellites could be bound by the MSL complex and engaged in LRCs specifically in embryos.

While in testes the mechanism underlying the LRCs between 1.688^3F^ satellites is likely different because they do not exhibit a pronounced binding of the MSL complex, but are enriched for the constitutive heterochromatin mark H3K9me2 (**Figure 3b**). This suggests that the LRCs could be mediated by AGO2 or other small RNA pathway genes that modulate the heterochromatin deposited at and nearby the 1.688^3F^ satellites(**Figure 3b**)(Deshpande and Meller, 2018). Since we (**Figure 1c**) and previous works(Witt *et al*., 2021) showed that DC is restricted to germline somatic cells and premeiotic cells in testes, we further examine the transcriptional activity of 1.688 satellites with the published single-cell RNA-seq data of *D. melanogaster* adult testis (Witt *et al*., 2019) to examine whether 1.688^3F^ satellites also have a cell type specific expression pattern. We find that only the X-linked 1.688 satellites, particularly the 1.688^3F^ satellites, have a high expression in the cyst cells (**Figure 3c**). All these results together establish an association between the stage and cell type specific patterns of expression, epigenetic status, and LRCs of 1.688^3F^ satellites that likely play a distinctive role from other 1.688 satellites and HAS/PinoX in coordinating the DC process.

The LRCs between the three classes of X-linked sequence elements could either function in parallel or compensate for each other’s function in different tissues or stages during the DC process. To test these two scenarios, we then cross checked the genomic locations and LRC frequencies of the three elements against the background of different types of DC genes that we defined above (**Figure 1d**). We find that PionXs and 1.688^3F^ satellites are predominantly located within SDC and FDC gene regions (including the flanking 5 kb regions), while HASs are predominantly located within CDCs. Moreover, the normalised LRC frequencies between the PionXs or the 1.688^3F^ satellites within the SDC/FDC genes are significantly (Wilcoxon test, P < 0.05) higher than those within the CDC genes, while those between HASs within the CDC genes are higher than those within the SDC/FDC genes (**Figure 3d, Supplementary Fig. 10a**). This result does not directly inform whether the elements or the genes are actually engaging in such discriminative LRCs. We next specifically look into the three classes of elements that do not overlap with any of the DC genes and they show a similar pattern: the ‘solo’ PionXs and 1.688^3F^ satellites much more frequently have LRCs with FDCs or/and SDCs than with CDCs, while the solo HASs show the opposite pattern (**Figure 3e**), with consistent patterns in both embryos and testes (**Supplementary Fig. 10b**). These results together indicate that HAS preferentially reside within and interact with CDCs, while PionXs and 1.688^3F^ satellites preferentially reside within and interact with FDCs/SDCs.

### Dosage compensation genes and sequence elements are differentially co-localized with TAD boundaries

Previous studies in Drosophila reported topologically associated domains (TADs) and contacts between the TAD boundaries (TABs) as an emerging developmental feature during the zygotic genome activation(Hug *et al*., 2017). These contacting TABs are maintained during later development, and are associated with RNA Polymerase II and enriched for housekeeping genes. CDC genes are found here to be enriched as well for housekeeping genes (**Figure 1**), and contacts between the TABs likely span long genomic distances. This inspired us to examine the associations between different types of DC genes, regulatory sequence elements of DC and 3D genomic features that may account for the LRC patterns that we observed above. We first divide the TAD into different chromatin states according to their combinatorial patterns of histone modifications (**Supplementary Fig. 11**). As expected, between 53% to 67% of the male-X linked TADs across development are characterised by overall high levels of MSL2 or H4K16ac bindings (the DC TAD) (**Supplementary Fig. 12a-b**) that encompass majorities of DC genes and regulatory elements (**Figure 4a, Supplementary Fig. 12c**). Substantial numbers (8% to 11%) of FDC genes, but not SDC genes are located in the Bivalent or Polycomb TADs that show high levels of H3K27me3 bindings, consistent with their gene level binding patterns (**Figure 1**).

**Figure 4:**
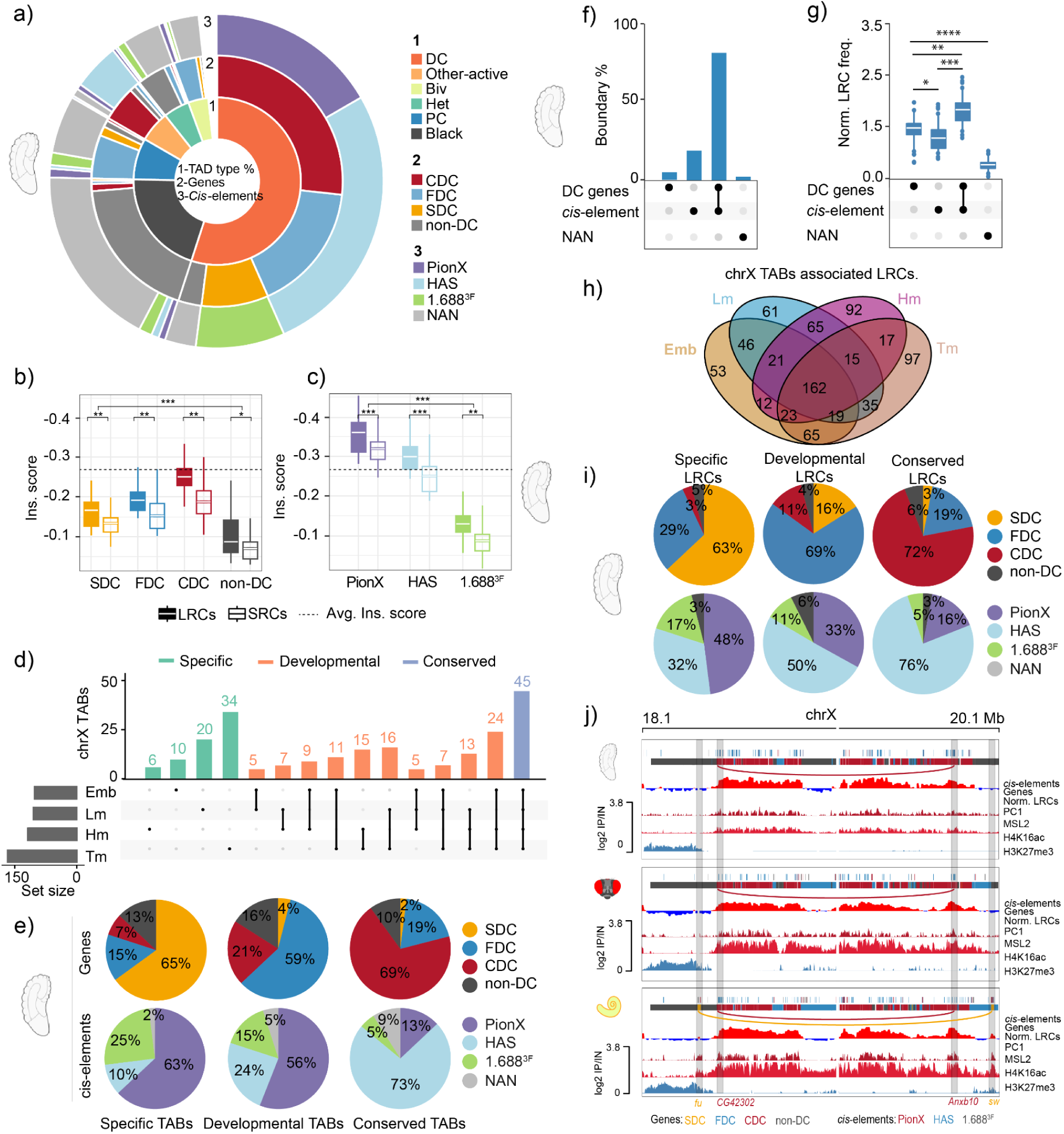
Dosage compensation genes and sequence elements are differentially co-localized with TAD boundaries. **(a)** The Circos plot illustrates the distribution of topologically associating domain (TAD) types along chrX, with the outer circles representing the percentage distribution of different DC gene classes and *cis*-elements, which are color-coded by category. **(b-c)** Box plots compare insulation scores at TAD boundaries across different gene classes and cis-elements, distinguishing between long-range contacts (LRCs) and short-range contacts (SRCs) with solid and empty boxes, respectively. Colors represent different gene and *cis*-element categories. Statistical significance was calculated using (Wilcoxon test, *P < 0.05, **P < 0.005, ***P < 0.0005). **(d)** An upset plot categorizes TAD boundaries into specific, developmental, and conserved types across developmental stages. **(e)** Pie charts detail the distribution of genes and *cis*-elements within TAB types, with each category represented by a unique color. **(f)** An upset plot displays the co-enrichment of chrX TAD boundaries with dosage compensated (DC) genes and *cis*-elements. **(g)** A boxplot compares the frequency of normalized long-range contacts across different TAD boundary classes. Statistical significance was calculated using (Wilcoxon test, *P < 0.05, **P < 0.005, ***P < 0.0005). **(h)** Venn diagram represents the overlap between TABs associated LRCs between different developmental stages and tissues. **(i)** Pie graph shows percentage distribution of DC genes (upper row) and regulatory sequences (below row) in embryo. **(j)** An illustrative example shows that genes under constitutive dosage compensation form stable LRCs, whereas genes compensated specifically in testes exhibit unique LRCs that are absent in other stage/tissue.

Genes with LRCs, regardless compensated or not, show a significantly (Wilcoxon test, *P*< 0.05) lower TAD insulation score, i.e., are more likely to co-localise with TABs than those without LRCs. Moreover, dosage compensated genes are more likely (Wilcoxon test, *P*< 0.05) than the uncompensated genes, CDC genes are more likely than the FDC/SDC genes, to co-localised with the TABs (**Figure 4b, Supplementary Fig. 12d**). Since majorities of HASs are encompassed within the CDC genes (**Figure 3d**), they are expectedly more likely than the 1.688^3F^ preferentially located in SDC genes to be co-localised with the TABs across all tissues/stages. PionXs only co-localise with the TABs in embryos and testis (**Figure 4c, Supplementary Fig. 12e**) where they exhibit LRCs with each other (**Figure 3**), suggesting formation of stage or tissue specific TAD structure whose boundaries encompass PinoXs and interact with each other. These patterns together suggested two types of LRCs associated with DC: the stable LRCs between CDC genes or HASs/PinoXs are mainly contacts between TABs; and the developmentally regulated LRCs between FDCs/SDCs are inter-TAD contacts that skip the TABs.

To test this hypothesis, we next divide the TABs into the conserved, developmental and specific TABs, after examining their presence/absence across development (**Figure 4d**). Stable LRCs predict an overrepresentation of CDC genes and HASs overlapped with the conserved TABs. This is indeed the case (**Figure 4e, Supplementary Fig. 13a**): around 70% of the DC genes or elements overlapped with conserved TABs are CDC genes and HASs, while nearly 60% of the DC genes or elements overlapped with developmental/specific TABs are FDC/SDC genes and PionXs/1.688^3F^ elements.

We next ask quantitatively how many DC genes and regulatory sequences are overlapped with the TABs. Across all stage/tissue that we sampled, we find between 72% and 78% of the TABs are overlapped with both DC genes and regulatory elements, but much fewer percentages of TAB overlap with only the DC gene (2% to 15%) or the regulatory element (5% to 11%) (**Figure 4f, Supplementary Fig. 13b**). Moreover, the frequencies of LRCs between TABs are significantly (Wilcoxon test, *P* < 0.05) higher in those encompassing both DC genes and regulatory elements than those containing either or none (**Figure 4g, Supplementary Fig. 13c**). After analyzing the higher LRC frequencies between TABs in those encompassing both DC genes and regulatory elements we divide these LRCs into specific, developmental, and conserved groups based on their presence or absence across developmental stages **(Figure 4h)**. The majority of CDC genes and HAS, ranging from 72% to 76%, colocalize with conserved LRs. In contrast, specific and developmental LRCs predominantly feature specific or facultative dosage compensated (SDC/FDC) genes and PionXs, as shown in (**Figure 4i and Supplementary Figure 14**). This suggests the formation of stage or tissue-specific topologically associating domain (TAD) structures, where boundaries include PionXs/1.6883F and facilitate interactions among them. Illustratively, CDC genes associated with HAS form stable LRCs, whereas SDC genes associated with PionXs/1.6883F display unique LRCs absent in other tissues (**Figure 4j**).

## Discussion

Dosage compensation is a regulatory process to restore the transcription balance of the entire chromosome or some individual genes between sex chromosomes and autosomes in the heterogametic sex. Here we focus on the first form that has been extensively studied for its mechanisms in genetic model organisms and ask how this process is finely tuned in the context of developmental variations and 3D genome architecture in Drosophila. We hypothesize that it is deleterious to unanimously change all the X-linked genes and provide evidence that around 30% of the genes are not compensated in one or more tissues or stages. These genes tend to have developmentally specific expression and functions and are enriched for polycomb epigenetic modifications. In addition, we uncover that the male chrX is characterised by a developmentally increasing long range contacts that are much weaker on autosomes and female chrXs. These LRCs are mainly between CDC genes and HAS elements that are overlapped with TABs that remain stable during the developmental course. While other developmentally regulated LRCs are preferentially associated with the PionX/1.688^3F^ sequence elements and developmentally regulated DC genes. These results together suggest that the male Drosophila chrX form two layers of topological architectures with a housekeeping architecture comprising stably interacting, transcribing, and compensated genes, and another with developmentally regulated ones involving different sets of genes and regulatory elements. An interesting pattern is that the 1.688^3F^ satellites were previously thought to not directly interact with the MSL complex, but here we showed their specific binding in the embryonic stage. Another interesting pattern is that the LRCs between TABs without the DC genes or sequence elements are much weaker, and also in certain tissue/stage the absence of TABs is associated with that of LRCs between PionXs. It remains to be tested in future whether the specific formation of TABs are in fact associated with that of LRCs and the responsible molecules for this fine regulation of DC.

## Methods

### Fly sample collection

*Drosophila melanogaster* stocks were maintained at 18-19 °C with a 12-hour light/dark cycle. We raised the flies on the Institute of Molecular Pathology (IMP) standard fly food with yeast in plastic bottles vials. For the larvae collection, we collected only the male larvae under a microscope, and for adult tissues like testes, and head samples, we sorted the virgin flies under the microscope and raised them on standard food for 3–5 days. For testes, dissected 3–5 days old virgin flies and collected 200-400 pairs of testes (samples) in cold testes extraction buffer (TEB) (10 mM HEPES, 100 mM NaCl,1xPBS, 1x protease inhibitors, 1 mM PMSF) while head tissue was extracted using glass beads along with liquid nitrogen.

### ChIP-seq and Cut and Run experiments

Tissue samples were homogenized in a buffer containing 140 mM NaCl, 1 mM EDTA, 10 mM HEPES, 0.1% Triton-X100, 1x protease inhibitors, and 1 mM PMSF. They were then cross-linked with 1% formaldehyde. The cross-linking was quenched by 125 mM glycine and 0.1% Triton-X-100. Samples were lysed in a solution of 50 mM Tris-HCl (pH 7.5), 10 mM EDTA, 1% SDS, protease inhibitors, and 1 mM PMSF and sonicated using an ultra-ultrasonicator. Fragmented chromatin was sedimented at high speed and resuspended in a cold nuclear lysis buffer containing 1x Protease inhibitor (halt), 1 mM PMSF, 10 mM Tris-HCl, 1 mM EDTA, 0.5% NP-40, 0.1% SDS, and 0.5% N-lauroylsarcosine. The chromatin was incubated with 2-3ul of the antibody (H3K36me3 (Abcam, #ab9050, Rabbit Polyclonal, 1 mg/mL, 3 µg/mL), H3K27me3 (Abcam, #ab6002, Mouse Monoclonal, 0.9 mg/mL, 2 µg/mL), H3K9me2 (Abcam, #ab1220, Mouse Monoclonal, 0.9 mg/mL, 2 µg/mL), and H4K16ac (Millipore, #07-329, Rabbit Polyclonal, 1 mg/mL, 2 µg/mL), MSL2 guinea pig (5ul per reaction, shared by Regnard lab/Becker lab) at 4 °C overnight. The chromatin-antibody complexes were then coupled with Pierce protein A/G magnetic beads and rotated at 4 °C for 2-3 hours. Beads were washed sequentially with RIPA, LiCl, and TE buffers. De-crosslinking was performed either for 6 hours or overnight at 65 °C using 4.5 µL of Proteinase K (20 mg/ml) and 5 µL of RNase A (0.5 mg/ml). The DNA was subsequently purified using a phenol-chloroform-isoamyl alcohol mixture (25:24:1 ratio). For H3K27ac and H3K36me3 markers in testes, we used the CUT and RUN technique by ref.(Skene and Henikoff, 2017). Tetis tissue homogenised and nuclei were extracted and incubated with an antibody in the dig-wash buffer (0.05% Digitonin, 2 mM EDTA,0.5 mM Spermidine, 10 mM PMSF,1x Protease Inhibitors) at 4 °C overnight on an end-to-end rotator. After centrifugation, the nuclei were washed and treated with pA-MNase and washed resuspended in dig-wash buffer and 2 µL of 100 mM CaCl2 was added, followed by incubation for 20-25 minutes at 0 °C. Next, the reaction was stopped by adding 150 µL of 2x stop buffer (200 mM NaCl, 20 mM EDTA, 4 mM EGTA, RNAse, 40ug/ml Glycogen) and samples were incubated at 37 °C for 10 minutes to release CUT&RUN fragments. Slightly spin the samples and the supernatant was transferred to a fresh tube, and the targeted chromatin was pelleted at high speed. DNA extraction was performed using a standard phenol-chloroform-isoamyl alcohol mixture. ChIP libraries were prepared with the New England Biolab’s NEBNext Ultra II DNA Library Prep Kit (E7645) and sequenced on the Illumina HiSeq platform by Novogene UK in 150PE mode.

For all the ChIP-seq datasets, we applied strict quality check and discarded any data that did not meet our following criteria. Our quality check includes first for each histone modification mark, we examine its binding distribution between active vs. inactive genes, coding vs. non-coding repetitive regions, and distribution along the gene body. We also manually examined many known genes’ gbrowser binding profiles across different tissues and stages regarding the expected broad or narrow binding patterns of respective marks. We performed deep sequencing (5 G) for each histone mark in each tissue, which provided high coverage and resolution, ensuring robust and reliable data. Quality control of raw reads was performed using FastQC and Illumina adapters were trimmed using trimmomatic(Bolger, Lohse and Usadel, 2014). Trimmed reads were mapped to the dm6 genome assembly using bowtie2(Langmead and Salzberg, 2012), with parameters (using the ‘XA:Z:’ and ‘SA:Z:’ tags created by bowtie2 mapping in the SAM file, we utilized SAMtools to filter out multi-mapped reads, allowing us to retain only uniquely mapped reads) set to permit only unique alignments. We identified target signal enrichment by calculating the standardized variance between the normalized immunoprecipitated signal and its matching normalized input coverage. We used MACS2(Zhang *et al*., 2008) to call call narrowpeaks using qvalue 0.01 and coverage files were generated with deepTools(Ramírez *et al*., 2014) bamCompare function using binsize of 10 bp (--bs 10 --minMappingQuality 10 --normalizeusing RPKM).

### Antibody generation and immunofluorescence analysis

Antibodies were raised in rabbits against peptides of the MSL2 CXC-domain. The affinity-purified antibodies were applied in immunofluorescence staining at the following dilutions: 1:1,000, anti-MSL2 (rabbit), 1:1000 anti-MSL2 (guinea pig,shared by Regnard lab/Becker lab) and 1:1000 for anti-H4K16ac (rabbit). Mounting of adult testes, anti-histone, and secondary antibodies was used. Early cyst cells were visualized by anti-TJ, and DNA was visualized with DAPI 33258 dye. Immunofluorescence was monitored using a Zeiss AxioPlan2 microscope equipped with appropriate fluorescence filters. Recorded images were processed with Image-J/Fiji. To first check the efficiency of our newly produced MSL2 antibody in *D. melanogaster*, the staining of the virgin adult testes with the affinity-purified MSL2 antibody (anti-MSL2) showed that the MSL2 protein transcribed in male testes. In contrast, the female ovaries showed weak or no anti-MSL2 staining. This confirms that the MSL2 protein is male-specific, as expected for a gene involved in dosage compensation. To this end, the staining of the *D. melanogaster* virgin male testes was performed in contrast with traffic jam (TJ) antibody (1:5000), which labels the testes early-cyst cell. As a control, the ovaries of *D. melanogaster* females were simultaneously stained. The results are shown in Figure 1. In *D. melanogaster* the anti-MSL2 was mainly stained in the early-cyst cells (labeled with TJ) and nicely overlapped with H4K16ac stained cells, which showed genes involved in dosage compensation.

### Defined MSL-complex binding sites HAS and PionX

The MSL-complex binding sites were defined as previously explained in three different studies (Ramírez *et al*., 2015), (Villa *et al*., 2016) and (Alekseyenko *et al*., 2012). CXC-dependent (PionX) and MSL2 induced high-affinity sites (HAS) were identified using the previously described definitions explained in (Ramírez *et al*., 2015) and (Villa *et al*., 2016). Briefly, HAS were previously identified as the localization of MSL-2 in *in-vivo* ChIP-seq Peaks and PionX sites are *in-vitro* DIP-seq MSL-2 sites (Villa *et al*., 2016) that lose binding upon deletion of its CXC-dependent domains (used previously published list). PionX sites in embryo, larvae, head and testes were identified using FIMO. The previously published PionX sites (Villa *et al*., 2016) were used to predict the PionX using FIMO against our MSL2 ChIP-seq peaks in corresponding stage/tissue. To run FIMO the following parameters were used: -motif 1 --verbosity 1 --thresh 1e-5 --qv-thresh --parse-genomic-coord.

### Identification of 1.688 satellite repeat copies

We identified 1.688 satellite repeat copies using BLASTn (/apps/ncbiblastplus/2.11.0/bin/blastn) -outfmt 6. Filter the blast hits that have at least 80% identity with minimum query coverage >= 70% and E-value >=1e-5 in order to prevent non-specific results. For 1.688^3F^ copy we make sure their relevant consensus sequence have at least one 100% mapped hit within 3F3 region. 1.688^1A^, 1.688^3c^, 1.688^7E^ and 1.688^7F^ copies were annotated and their consensus sequences are labeled as 1.688^non-3F^ in this study. We separated the X-linked satellite repeat copies based on their sequence similarities and the total repeats higher on chrX than autosomes and the rest of the consensus sequence copies were labelled as 1.688^non-X^. To further prevent any overlaps between these different classes of satellite repeats, we used bedtools intersect to get non-overlapped regions (unique hits) for each category.

### Transcriptomic analysis

Total RNA alignment was performed using RSEM (Version_1.3.1)(Li and Dewey, 2011) against the reference transcript sequences and gene annotation (reference: dm6.gtf) using bowtie2 with default parameters. Reads were counted per transcript and summed for each gene using the “rsem-calculate-expression” function in the RSEM-package. Male vs female log2-fold change gene expression levels were calculated using the DESeq2 (Love, Huber and Anders, 2014)package. To test whether there are differences in gene expression level during development, TAU score was calculated (higher the TAU-value means more stage/tissue specificity). Differentially expressed genes are listed in Supplementary Data.

### Small RNA-seq analysis

Small RNA-seq reads were downloaded from NCBI (listed in supplementary Table) and reanalyzed in our study. Adapters were trimmed using trimmomatic and trimmed reads were then mapped to the reference genome (dm6) using Bowtie (Langmead, 2010). To guarantee that, reported alignments are unique –m value was set to 1 (-m 1). To count the reads that uniquely mapped to each Repeat feature type, a customized python script was used (https://github.com/LarracuenteLab/Dmelanogaster_satDNA_regulation (Wei *et al*., 2021), htseq_bam_count_proportional.py). We further compare the size distribution and relative nucleotide bias at positions along each satDNA by extracting reads mapped to the satDNA of interest (1.688 satDNA) using a customized python script (https://github.com/LarracuenteLab/Dmelanogaster_satDNA_regulation (Wei *et al*., 2021), extract_sequence_by_feature_gff.py). All plotting analysis was done in R(*A Language and Environment for Statistical ## Computing_. R Foundation for Statistical Computing*, no date) (using Bioconductor package).

### Definition of DC genes

To classify male chrX genes as DC or non-DC genes, we only considered the active genes that have (FPKM >=1). Getting benefited by the differential distribution properties of H4K16ac and MSL2 along chrX vs autosomes, where along chrX its distribution is towards the 3-prime end and along autosomes, it is enriched towards the 5-prime end. We plotted their distribution chrX vs auto. and decided a cutoff where these two distributions were separated and using this cutoff value we successfully defined our DC and non-DC genes. Finally, our defined DC genes are either bound by H4K16ac or MSL2. We further separated constitutive dosage compensated (CDC) genes, facultative dosage compensated (FDC) genes and specific dosage compensated (SDC) genes using Intervene (Khan and Mathelier, 2017) (Version_0.6.5).

### Gene Ontology enrichment of biological processes

Gene Ontology Biological Processes (GOBP) enrichment was performed using Metascape(Zhou *et al*., 2019) for SDC, FDC and CDC genes as a target.

### Hi-C

*D. melanogaster* sexed Hi-C data sets were produced for 3^rd^ instar larvae, head, Ovary and Testes using DpnII as restriction enzyme was carried out as described by(Belton *et al*., 2012). While 16-18h old sex-sorted embryos Hi-C data sets were used from NCBI under accession (GSE94115). Briefly, Cells were cross-linked using 1% formaldehyde, and the reaction was subsequently quenched with 200 mM glycine to terminate cross-linking. The fixed cells were then used for Hi-C library preparation following the manufacturer’s protocol for Proximo Hi-C Kits v4.0 (Phase Genomics). The final amplified libraries were sequenced on the Illumina HiSeq X Ten platform (San Diego, CA, USA) using 150 bp paired-end (PE) mode.

### Hi-C data bioinformatics analysis

Sequencing reads were mapped to the Drosophila reference genome assembly considering only chromosomes 2, 3 and X. We excluded chrY and chr4 from the analysis and the reason is that either they are mainly heterochromatic or very short in size. We first used Hi-C pro utilities to Identify the genomic locations of each DpnII restriction site, the following command was used (digest_genome.py -r DpnII). Non-digested, self-ligated fragments were removed and reads were initially counted for each valid DpnII fragment to generate a fragment-level Hi-C object. Restriction fragment level Hi-C objects were then merged into bins of equal size at different resolutions, including 5, 10, 25 and 100 kb resolutions. Finally, the binned matrices were normalized using the iterative correction method from(Lyu, Liu and Wu, 2020). Moreover, to remove non-informative read pairs we excluded the first two diagonals during normalization (contacts at distances 0 or 1 bin).

For ICE normalization of Hi-C contact maps, we used the iterative_mapping module (Imakaev *et al*., 2012) (iced version 0.5.2) in HiC-Pro for aligning reads to the reference genome, making the assumption of equal visibility of each fragment. The following filtering parameters were adopted in ICE pipeline: --filter_low_counts_perc 0.02 and --filter_high_counts_perc 0, we used --max_iter 100, --eps 0.1 and --remove-all-zeros-loci, --output-bias 1, --verbose 1. We also filtered for duplicated or non-informative read pairs associated with PCR artifacts that were discarded during normalization. Only valid pairs involving two different restriction fragments were used to build the ICE–normalized contact matrices. The biological replicated from sex-sorted embryos Hi-C were mapped separately and then merged into their corresponding samples.

### Hi-C data downstream analysis

To further analyze these ICE-normalized matrices, we convert them to h5 format using ‘hicConvertFormat’ from HiCExplorer (Wolff *et al*., 2020) (version 3.6). Hi-C matrix visualization was performed using ‘hicPlotMatrix’ or ‘pyGenomeTracks’ for the specified regions.

TADs were called using ‘hicCallTADs’ and the following parameters were adopted ‘--correctForMultipleTesting fdr --numberOfProcessors 30 --minBoundaryDistance 5000 --thresholdComparisons 0.01 --delta 0.05 --step 5000’. That way, we obtained the TAD boundary positions and genome-wide TAD insulation scores. Conserved and non-conserved TABs between different developmental stages (embryo and larvae) and adult tissues (head and testes) were calculated (by extending the TABs by 2 kb) using bedTools intersect (version v2.29.2) and the minimum overlap cutoff was set to 50% (-f 0.5).

### TAD boundary conservation during development

To separate the constitutive or facultative TABs, we checked their overlap using bedTools intersect –f 0.7 –r. Next, for each domain border (constitutive or facultative), we considered a window with a size of up to 5 bins (50 kb) on both sides. If such windows overlap for any pair of neighboring domain boundaries, they are shortened as housekeeping or conserved boundaries intervening regions. This was an important point as it avoids overestimating the association of any boundary class to genomic features while allowing at the same time a definition of boundaries at fine-scale (i.e. small domains). Tissue or stage-specific TABs are defined as they only differentially appear in one tissue or stage but disappear from others.

### Defining chromatin state domains

ChromHMM(Ernst and Kellis, 2017) was used to identify chromatin states across the genome. We selected a 15-state model, as it effectively captured chromatin states that correspond to known biological processes and chromatin configurations, ensuring both depth and clarity in the results while adequately representing all possible combinations.To begin, we prepared and provided annotation files that included genomic features such as genes, transcription start sites (TSS), transcription end sites (TES), introns, exons, and transposable elements (TEs).

Additionally, we incorporated ChIP-seq histone post-translational modifications (HPTMs) along with the input control, using a single-cell type option. We then performed binarization at a 200 bp resolution, identifying regions of significant enrichment. Next, we combined the enrichment profiles of each chromosome from the previous step and trained the program to generate models with different numbers of chromatin states. Through this iterative process, we optimized the classification of chromatin states, ensuring a comprehensive representation of epigenetic landscapes across the genome.

### Computing Hi-C signal decay

hicPlotDistVsCount function from HiCExplorer was used to compute the interaction decay differences at close or long-range distances between dosage-compensated chrX and autosomes. For this, corrected Hi-C matrices of 50kb resolution were used to compute the Hi-C counts (along y-axis) vs distance (along X-axis) for the sexed Hi-C matrices separately, parameters --maxdepth 20000000 and --perchr.

### Compare matrices between sexes

To check detectable differences in the Hi-C interaction profiles in male dosage compensated chrX compared to female chrX. We compared male over female log2-fold ratio of contact frequencies of normalized Hi-C matrices binned at 25 kb resolution is shown for a representative 5Mb region of chrX. We performed fold-change comparisons using FAN-C (Kruse, Hug and Vaquerizas, 2020) (fig 1a). commands were used: fanc compare -c fold-change -l -Z –I –tmp to create fold-change matrices and then plot using command: fancplot X:5mb-10mb -p triangular -m 2.5mb -c RdBu_r -vmin -5 -vmax 5.

### Principal Component Analysis (PCA) saddle plot

The first eigenvector (PC1) corresponding to active (A) and inactive (B) compartments, was computed using FANC(Kruse, Hug and Vaquerizas, 2020). Corrected Hi-C matrices of 25 kb resolution were used to call compartments. To switch the orientation of PC1 values, where positive values correspond to the active compartment (A) and negative values correspond to the inactive compartment (B), we used GC contents. In the end, we verified the PC1 orientation for each chromosome to overlay with active and inactive histone modification mark ChIP-seq data. Saddle plot shows preferential contacts of active vs active (AA) in red and inactive vs inactive (BB) regions in blue color, and the bar plot on top shows the cutoffs used for binning regions by the corresponding EV entry magnitude.

### Aggregated Hi-C contacts

We used a common approach to summarize the average pairwise Hi-C contacts between a set of selected genomic loci. hicAggregateContacts from HicExplorer with corrected Hi-C matrices and settings “*--numberOfBins 60 --vMin 1 --vMax 2 --range 250000:3000000 --plotType 3d –avgType mean –chromosomes X –transform obs/exp*” to plot aggregated pairwise Hi-C contacts between a set of high-confidence MSL2 binding sites (HAS) on the X chromosome in matrices of resolution 25kb. We also checked the pairwise HAS contacts within the autosome (chr2L) as control.Enriched Hi-C contacts between HAS-HAS, pPionX-pPionX or HAS-pPionX along the X chromosome were visualized using aggregate plots as described above. The window of size n is created on the row and column indices, i and j respectively. In our case, the binning resolution of the Hi-C matrix was 5 kb and n = 60. Therefore, the examined window was of size ±150 or 300 kb surrounding any given pair of loci.

